# Binocular integration of retinal motion information underlies optic flow processing by the cortex

**DOI:** 10.1101/2020.10.16.342402

**Authors:** Rune N. Rasmussen, Akihiro Matsumoto, Simon Arvin, Keisuke Yonehara

## Abstract

Locomotion creates various patterns of optic flow on the retina, which provide the observer with information about their movement relative to the environment. However, it is unclear how these optic flow patterns are encoded by the cortex. Here we use two-photon calcium imaging in awake mice to systematically map monocular and binocular responses to horizontal motion in four areas of the visual cortex. We find that neurons selective to translational or rotational optic flow are abundant in higher visual areas, whereas neurons suppressed by binocular motion are more common in the primary visual cortex. Disruption of retinal direction selectivity in *Frmd7* mutant mice reduces the number of translation-selective neurons in the primary visual cortex, and translation- and rotation-selective neurons as well as binocular direction-selective neurons in the rostrolateral and anterior visual cortex, blurring the functional distinction between primary and higher visual areas. Thus, optic flow representations in specific areas of the visual cortex rely on binocular integration of motion information from the retina.

## Introduction

The action of moving through an environment produces patterns of visual motion, known as optic flow, on the retina, which animals rely on to guide their behavior. Animal locomotion is largely described by a combination of forward-backward movements and left-right turning. Forward and backward movements induce translational optic flow (nasal-to-temporal or temporal-to-nasal motion in both eyes, respectively) whereas turning induces rotational optic flow (nasal-to-temporal motion in one eye and temporal-to-nasal in the other) (Fig. 1a). However, despite the increasing use of mice to study vision, it is unknown how these distinct optic flow patterns are encoded by the rodent cortex.

**Fig. 1.**
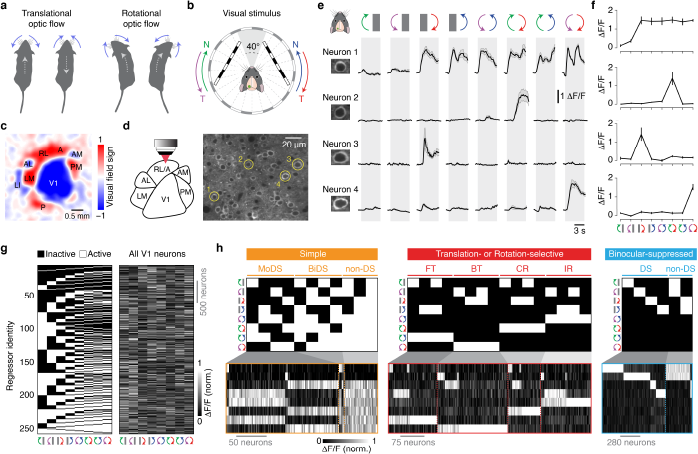
Discrete neuronal responses to motion stimuli can be imaged in the visual cortex of awake mice. **a**, Diagram illustrating optic flow patterns induced by self-motion. Forward and backward movements induce translational optic flow (left), and leftward and rightward turns induce rotational optic flow (right). Blue arrows indicate the dominant apparent motions in the visual space surrounding the mouse; gray dotted arrows indicate direction of locomotion. **b**, Diagram of the visual stimulus setup. Spherically-corrected gratings moved in either nasal (N) or temporal (T) directions (10 or 40 °/s with 0.03 cycles/°). The stimulus was not displayed in the binocular visual field (frontal 40°) to ensure stimulation of only the monocular visual fields. Imaging was performed in the visual cortex of the left hemisphere. **c**, Visual field sign map obtained with intrinsic signal optical imaging showing the location of visual cortical areas. **d**, Left: two-photon imaging was performed from identified visual cortical areas. Right: example image of GCaMP6f-expressing neurons in layer 2/3 of V1. **e**, Example trial-averaged fluorescence intensity (ΔF/F) time courses for the neurons highlighted in (**d**) in response to monocular and binocular motion. Error bars are mean ± s.e.m. **f**, Tuning curves of the neurons in (**e**). Error bars are mean ± s.e.m. **g**, Left: map of all 256 regressors. Right: response matrix of the tuning curves for all consistently-responsive V1 neurons. **h**, Regressor profiles and tuning curves for V1 neurons assigned to functional groups within the simple, translation- or rotation-selective, and binocular-suppressed response classes. MoDS, monocular DS; BiDS, binocular DS; FT, forward translational; BT, backward translational; CR, contraversive rotational; IR, ipsiversive rotational.

An extensive body of research has shown that neurons residing in brain areas involved in optic flow processing have complex receptive fields, often receive binocular inputs, and respond to both translational and rotational optic flow stimuli. Examples include the fly lobula plate (involved in course control)^1,2^, the zebrafish pretectal nuclei^3,4^, the avian and mammalian accessory optic system (involved in gaze stabilization)^5,6^, and both the dorsomedial region of the medial superior temporal area and posterior parietal cortex (PPC) of monkeys (involved in spatial navigation)^7-9^. The mouse visual cortex contains a primary visual cortex (V1) and more than a dozen distinct higher visual areas (HVAs), each with unique sensitivities to visual features^10,11^. The V1 receives retinal inputs via the lateral geniculate nucleus and distributes functionally specialized signals to different HVAs^12-14^. Based on their anatomy, multi-sensory processing, and roles in spatial navigation, the rostrolateral (RL), anterior (A), and anteromedial (AM) HVAs are considered part of the PPC in mice^15–18^, raising the possibility that they contain neurons sensitive to binocular optic flow.

In rodents, visual motion computations are not exclusive to the cortex, and start in the retina. The retina contains mosaic arrangements of direction-selective (DS) cells that preferentially respond to motion in one of the four cardinal directions (nasal, temporal, dorsal, and ventral)^19–21^. These cells fall into two canonical classes: ON DS cells (which project to the nuclei of the accessory optic system and mediate the optokinetic reflex) and ON-OFF DS cells (which project to the lateral geniculate nucleus and the superior colliculus)^19,21–23^. Interestingly, disruption of horizontal direction selectivity in the retina impairs monocular motion responses in layer 2/3 of V1 and the RL area, but not in the posteromedial (PM) area nor layer 4 of V1, suggestive of a segregated cortical pathway for processing signals originating from retinal ON-OFF DS cells^12,13,24,25^. An intriguing hypothesis is that information from ON-OFF DS cells in the left and right eyes is systematically integrated in the cortex to create areas with distinct sensitivity to translational and rotational optic flow patterns^21,26^. However, this has yet to be experimentally tested, and the cortical areas that might combine optic flow information from the left and right eyes remain unknown.

Here we systematically map the responses of individual neurons across the visual cortex using two-photon calcium imaging during monocular and binocular optic flow stimulation in awake mice. We test the contribution of retinal DS cells using *Frmd7* mutant (*Frmd7^tm^*) mice, in which retinal horizontal direction selectivity is disrupted^13,24,27^. Our data demonstrate that the mouse visual cortex contains an abundance of neurons that encode translational or rotational optic flow. Furthermore, our results suggest that information from retinal DS cells in each eye is integrated in the cortex as early as in V1, where it establishes response selectivity to backward translational optic flow, but that binocular retinal DS signaling for establishing selectivity to rotational optic flow is first integrated in the higher areas RL and A. Conversely, our finding that neurons suppressed by binocular motion and robustly activated by monocular motion are not impaired by disruption of retinal direction selectivity, supports the hypothesis that retinal ON-OFF DS cell mosaics are specialized for detecting translational and rotational optic flow rather than local motion^26^.

## Results

### Discrete neuronal responses to monocular and binocular motion stimuli can be imaged in the visual cortex of awake mice

To identify individual areas of mouse visual cortex, we used intrinsic signal optical imaging^13,28^. We first generated visual field sign maps from retinotopic maps, allowing us to identify V1 as well as the higher areas RL, A, AM, and PM (Fig. 1c and Extended Data Fig. 1). We chose to combine areas RL and A (RL/A), as these areas could not be not clearly distinguished from each other in our dataset^10,29^. For binocular animals to reliably detect different optic flow patterns, the brain must integrate motion signals from each eye. We therefore investigated the neuronal responses underlying binocular optic flow processing by presenting moving gratings to mice using a stimulus protocol that tests the repertoire of horizontal motions^3^. The eight stimulus conditions in the protocol were generated by presenting gratings moving in a nasal or temporal direction to one eye at a time, and then to both eyes to simulate the rotational (ipsiversive and contraversive) or translational (forward and backward) optic flow that the mouse would experience during locomotion (Fig. 1a,b and Supplementary Video 1; see Methods). To unambiguously probe the interaction of left and right retinal information in the cortex, the stimuli were presented only to the monocular visual fields and not to the frontal binocular visual field (Fig. 1b). Our stimulus protocol did not trigger the optokinetic reflex (Extended Data Fig. 2), likely due to the use of a low spatial frequency^30^.

**Fig. 2.**
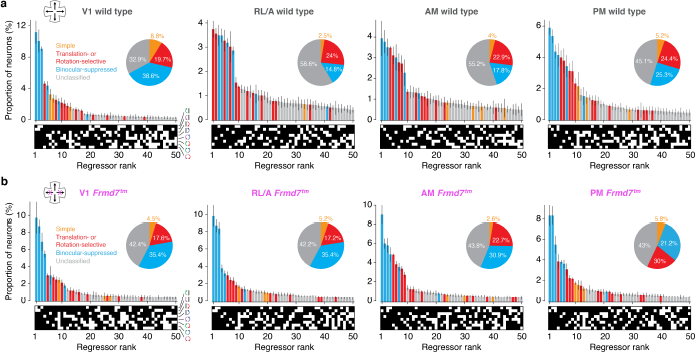
The RL/A area of the visual cortex is enriched with optic flow-selective neurons in wild-type mice. **a,b**, Ranked distribution of the 50 most abundant regressor profiles and response classes in the V1, RL/A, AM, and PM areas of wild-type mice (**a**) and *Frmd7^tm^* mice with disrupted retinal direction selectivity along the horizontal axis (**b**). Error bars are mean ± s.e.m. Inset: proportion of all neurons in the response classes.

The tuning properties of individual layer 2/3 neurons were characterized in awake mice by transfecting cortical neurons with the genetically encoded calcium sensor GCaMP6f and measuring changes in two-photon fluorescence during stimulus presentation (Fig. 1d,e). A typical field of view contained ~100–150 neurons, and somatic calcium responses showed diverse, but consistent, patterns, depending on the eye being stimulated and the direction of motion (Fig. 1e). Tuning curves for individual neurons were generated by plotting trial-averaged fluorescence changes as a function of stimulus conditions (Fig. 1f). We systematically classified neurons into distinct types according to their tuning curves using regressor-correlation analysis^3^. First, we generated a regressor map consisting of all possible all-or-none response combinations to the eight stimulus conditions, which resulted in 256 profiles (Fig. 1g; see Methods). Next, the tuning curve for each neuron was assigned to the regressor with the highest correlation (Extended Data Fig. 3). All tuning curves had high correlations with their assigned regressor (mean correlation coefficient, 0.91 ± 0.05, *n* = 26712 neurons from 17 mice). These data confirm that we can reliably elicit responses to monocular and binocular motion stimuli in the visual cortex of awake mice and also robustly classify neurons into discrete response types.

**Fig. 3.**
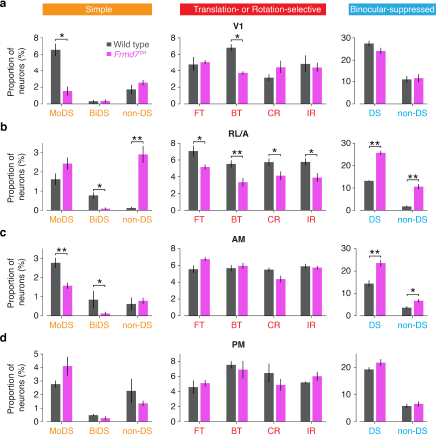
Retinal direction selectivity contributes to optic flow-selective responses in an area-specific manner. **a,b,c,d** Proportion of V1 (**a**), RL/A (**b**), AM (**c**), and PM (**d**) neurons in simple, translation- or rotation-selective, and binocular-suppressed functional groups for wild-type and *Frmd7^tm^* mice (*P < 0.05, * *P < 0.01, two-way ANOVA with two-sided Mann-Whitney *U* tests for post hoc comparisons, *n* = 4 mice for V1 and PM and *n* = 5 mice for RL/A and AM). Error bars are mean ± s.e.m.

### The RL/A area of the visual cortex is enriched with optic flow-selective neurons

We sought to investigate the response specificity of visual cortex neurons by sampling thousands of consistently-responsive neurons in multiple areas of the visual cortex of nine mice (3010 in V1, 4165 in RL/A, 4006 in AM, and 3059 in PM; Supplementary Table 1) and assigning them to regressors (Fig. 1g and Extended Data Fig. 4). To characterize the monocular and binocular optic flow coding properties of these neurons, we initially focused on three response classes: simple, translation- or rotation-selective^3^, and binocular-suppressed. The simple class comprised three groups that were characterized by their direction selectivity: monocular DS, binocular DS, and non-DS neurons (Fig. 1h). Translation- and rotation-selective neurons comprised four groups that were characterized by their response selectivity to either forward translational, backward translational, contraversive rotational, or ipsiversive rotational optic flow (Fig. 1a,h). Binocular-suppressed neurons were characterized by a suppressed response during binocular motion stimulation and were further divided according to their DS or non-DS responses to monocular motion (Fig. 1h; see Methods).

**Fig. 4.**
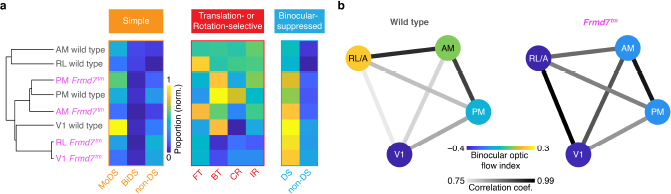
Retinal direction selectivity establishes functional segregation between V1 and RL/A. **a**, Left: Hierarchy showing similarity in proportion of functional groups between visual areas in wild-type and *Frmd7^tm^* mice. Right: mean proportion of neurons in simple, translation- or rotation-selective, and binocular-suppressed functional groups between visual areas in wild-type and *Frmd7^tm^* mice, sorted according to the similarity hierarchy (left). **b**, Diagram of the binocular optic flow index for each visual area, and the correlation in functional group proportions between areas, in wild-type and *Frmd7^tm^* mice.

For all visual cortical areas, we counted neurons assigned to each regressor and ranked regressors according to their frequency (Fig. 2a and Extended Data Fig. 3). Interestingly, in contrast to previous work in zebrafish^3^, the most abundant neurons in V1 were binocular-suppressed neurons, which have been described in the primate V1^31^. These neurons constituted as much as 38.6% of V1 neurons and 50% of the 10 most frequent regressors (Fig. 2a). In contrast, simple and translation- or rotation-selective neurons constituted only 8.8% and 19.7% of all responsive neurons, respectively (Fig. 2a). Neurons that could not be assigned to these three classes were considered unclassified and not investigated further.

The abundance of neuronal classes was different in the HVAs (Fig. 2a). Translation- or rotation-selective neurons were the most abundant response class in the RL/A area (24% of neurons, corresponding to 50% of the 10 most frequent regressors), whereas simple and binocular-suppressed neurons comprised only 2.5% and 14.8% of neurons, respectively. In area AM, translation- or rotation-selective neurons were again abundant and simple neurons sparse (22.9% and 4% of neurons, respectively), but there was a higher proportion of binocular-suppressed neurons than in the RL/A area (17.8%). The PM area was characterized by an equal proportion of translation- or rotation-selective and binocular-suppressed neurons, constituting 24.4% and 25.3% of neurons, respectively (Fig. 2a).

These data establish that different areas of mouse visual cortex contain distinct distributions of monocular and binocular optic flow-encoding neurons. In particular, the RL/A area is enriched with neurons encoding translational and rotational optic flow, whereas V1 is enriched with neurons activated by monocular motion but suppressed by binocular motion.

### Retinal direction selectivity contributes to binocular optic flow processing in V1 and RL/A

To determine whether retinal direction selectivity contributes to the processing of optic flow in the visual cortex, we repeated our neuronal mapping in *Frmd7^tm^* mice, which lack horizontal direction selectivity in the retina^13,24,25,27^. Consistently-responsive neurons were sampled in different areas of the visual cortex of eight mice (2925 in V1, 3125 in RL/A, 3375 in AM, and 3047 in PM; Supplementary Table 1). This revealed a difference in the overall distribution of response classes in certain areas of *Frmd7^tm^* mice compared to wild-type mice (Fig. 2a,b), which prompted us to examine the effects of direction selectivity on the proportions of monocular- and binocular-responsive neurons in each response class (Fig. 3a-d). In V1, the proportions of monocular DS and backward translation-selective neurons were reduced in *Frmd7^tm^* mice (Fig. 3a). More strikingly, all groups of translation- or rotation-selective neurons, as well as binocular DS neurons, were reduced in the RL/A area of *Frmd7^tm^* mice (Fig. 3b). In the AM area, only monocular and binocular DS neurons were reduced (Fig. 3c). The proportion of DS and non-DS binocular-suppressed neurons was increased in both RL/A and AM areas of *Frmd7^tm^* mice (Fig. 3b,c). Finally, none of the nine functional groups were significantly altered in the PM area of *Frmd7^tm^* mice (Fig. 3d), underscoring previous work showing that motion processing in the PM area is independent of retinal DS signaling^13^.

Together, these data show that simple and translation- or rotation-selective responses, but not binocular-suppressed responses, are impaired by disrupting retinal direction selectivity. Furthermore, we conclude that retinal direction selectivity contributes to binocular optic flow responses in the V1 and RL/A areas of the visual cortex.

### Retinal direction selectivity establishes functional segregation between V1 and RL/A

Individual HVAs form distinct subnetworks, each of which represents a different information stream^10,32,33^. We sought to find out how visual cortical areas are functionally organized with respect to their composition of optic flow-sensitive neurons, and if retinal direction selectivity is involved in creating such an organization. To probe this, we used the mean proportion of neurons in our nine functional groups to create an optic flow fingerprint for each visual area in wild-type and *Frmd7^tm^* mice, then performed hierarchical clustering (Fig. 4a) and correlation analyses (Fig. 4b; see Methods).

Hierarchical segregation (Fig. 4a) and a weak correlation between optic flow representations (mean correlation coefficient, 0.89 ± 0.03; Fig. 4b) were evident between the cortical areas of wild-type mice. In particular, V1 was separated from the RL/A, AM, and PM areas, suggesting functional specialization between V1 and the HVAs^10^. In addition, the PPC areas (RL/A and AM) branched early from V1 and PM, indicating that the PPC has a distinct role in optic flow processing (Fig. 4a). In contrast, there was little hierarchical segregation, and more correlated optic flow representations, between visual areas in *Frmd7^tm^* mice (mean correlation coefficient, 0.96 ± 0.007; Fig. 4a,b). Notably, optic flow responses in area RL/A were remarkably similar to those in V1 in *Frmd7^tm^* mice (correlation coefficient, 0.75 and 0.99, for wild-type and *Frmd7^tm^* mice, respectively; *P* < 0.01, Fischer’s transformation, *n* = 9 proportion values; Fig. 4b), abolishing any functional segregation between these areas. In contrast, the PM area of both wild-type and *Frmd7^tm^* mice appeared on the same branch (Fig. 4a), supporting the notion that motion processing in this area is independent of retinal direction selectivity.

To further investigate area specialization, we assessed the proportion of monocular- versus binocular-driven functional groups within each visual area and quantified the relationship with a selectivity index (Fig. 4b; see Methods). In wild-type mice, the bias towards monocular or binocular motion differed between visual areas to the extent that RL/A emerged as a specialized area for binocular optic flow processing (binocular optic flow index, −0.39 for V1, 0.21 for RL/A, 0.059 for AM, and −0.11 for PM; Fig. 4b). In contrast, this functional diversity was absent in *Frmd7^tm^* mice, and monocular-driven neurons were overrepresented across the visual areas (binocular optic flow index, –0.38 for V1, –0.44 for RL/A, –0.18 for AM, and –0.19 for PM; Fig. 4b).

From these data we conclude that retinal direction selectivity contributes to functional segregation and response specialization between the different areas of the visual cortex in wild-type mice. The most striking effect of retinal direction selectivity disruption in *Frmd7^tm^* mice is the transformation of optic flow responses in the RL/A area into V1-like responses, indicating a specific role for the RL/A area in binocular integration of motion information originating from retinal DS cells.

## Discussion

Our study provides four major insights into the functional organization of optic flow processing in the visual system of mice. First, translation- and rotation-selective neurons are abundant in areas RL/A, AM, and PM, whereas neurons suppressed by binocular motion are common in V1. Second, translation-selective neurons in V1, and translation- and rotation-selective neurons in the RL/A but not AM and PM areas, rely on direction selectivity that is computed in the retina. Third, binocular-suppressed neurons, which would be efficiently activated by monocularly-restricted “local” motion but suppressed by self-motion-induced optic flow, do not rely on retinal direction selectivity. Fourth, retinal direction selectivity contributes to the functional segregation of optic flow responses between V1 and RL/A. Our results, therefore, demonstrate a causal link between retinal motion computations and optic flow representations in specific areas of the visual cortex. Furthermore, they establish a critical role for retinal direction selectivity in the cortical processing of whole-field optic flow, rather than local motion, thereby answering a previously proposed hypothesis^26^.

The altered optic flow representations in *Frmd7^tm^* mice imply potential functional circuits to link retinal horizontal DS cells and cortical layer 2/3 neurons with distinct optic flow response preferences (Fig. 5). Our results suggest that information from retinal DS cells, tuned to motion in either the nasal or temporal direction, is propagated to layer 2/3 of the contralateral V1, where it contributes to establishing monocular DS responses tuned to horizontal motion. In turn, a fraction of backward translation-selective responses in V1 are likely synthesized from these monocular DS inputs, converging from V1 in both hemispheres via interhemispherically-projecting neurons^34^. In addition, a fraction of rotation-selective responses in area RL/A are likely synthesized from monocular nasal- and temporal-motion-preferring DS inputs converging from V1 in the same and opposite hemisphere, respectively. These hypotheses could be tested by functionally characterizing the pre-synaptic network of individual translation- or rotation-selective neurons using rabies virus-based trans-synaptic tracing^35,36^. Our data also suggest that translation- and rotation-selective neurons in V1 and RL/A are suppressed by visual motion in non-preferred directions on either retina (Fig. 5). Such response suppression could be mediated by inhibitory monocular DS neurons or inhibitory interneurons activated by excitatory monocular DS neurons. Future studies could clarify this by genetically assigning imaged neurons into excitatory and inhibitory cell types.

**Fig 5.**
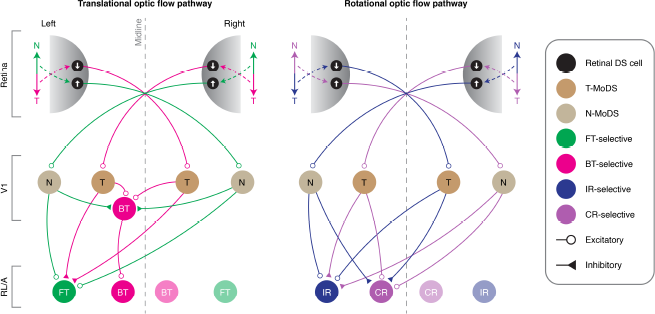
Proposed circuit model for translational and rotational optic flow processing. Left: FT optic flow activates nasal motion-preferring DS cells in the left and right retinas, mediating activity in nasal (N) motion-preferring MoDS (N-MoDS) neurons in V1 of both hemispheres, and subsequently their combination in FT-selective neurons in area RL/A. Activity in N-MoDS neurons also inhibits BT-selective neurons. BT optic flow activates temporal (T) motion-preferring DS cells in the left and right retinas, mediating activity in temporal motion-preferring MoDS (T-MoDS) neurons in V1 of both hemispheres, and subsequently their combination in BT-selective neurons in V1 and RL/A. Activity in T-MoDS neurons also inhibits FT-selective neurons. Right: IR optic flow activates temporal and nasal motion-preferring DS cells in the left and right retinas, respectively, mediating activity in N- and T-MoDS neurons in V1 of the left and right hemispheres, respectively. The signals from these V1 neurons, in turn, combine at IR-selective neurons in RL/A of the left hemisphere, and their activity inhibits CR-selective neurons in the left hemisphere. CR optic flow activates nasal and temporal motion-preferring DS cells in the left and right retinas, respectively, mediating activity in T- and N-MoDS neurons in V1 of the left and right hemispheres, respectively. The signals from these V1 neurons, in turn, combine at CR-selective neurons in RL/A of the left hemisphere, and their activity inhibits IR-selective neurons of the left hemisphere. The wiring diagram is expected to be mirror-symmetric in relation to the midline.

Our results also offer insights into the cortical pathways that process visual motion independently of direction selectivity computed in the retina. Our analyses reveal that neuronal responses suppressed by binocular motion are common in V1 and HVAs, and that these do not rely on retinal direction selectivity. This suggests that the V1 circuitry associated with binocular-suppressed neurons is functionally segregated from the circuitry processing retinal direction selectivity^12,13,24,25^. Interestingly, the majority of binocular-suppressed neurons in V1 had a preference for motion in the ipsilateral eye (Fig. 1h), suggesting that these neurons may combine the following two distinct types of input: 1) DS or non-DS excitatory inputs originating from non-DS cells in the ipsilateral eye, via interhemispherically-projecting neurons in the contralateral V1; and 2) non-DS inhibitory inputs driven by the activity of the contralateral eye. Our analyses also detected retinal DS cell-independent binocular optic flow responses in layer 2/3 of the visual cortex (Fig. 3b). Prior work in monkeys showed that binocular-suppressed and binocular-facilitated responses of monocular V1 neurons can be observed in the main visual input layer (layer 4)^31^. In mice, one form of *de novo* direction selectivity emerges in layer 4^37^. Hence, it is plausible that retinal direction selectivity-independent forms of binocular-suppressed and binocular-facilitated DS responses may arise in layer 4 from binocular interactions of DS signals originating from cortically-computed direction selectivity. This idea is consonant with a previous study in mice demonstrating that layer 4 neurons in V1 generate directionally-tuned responses independent of inputs from retinal DS cells^13^.

Accumulating evidence suggests that areas RL and A are part of the PPC in mice^15–18^ – a key nexus of sensorimotor integration that is involved in decision-making during spatial navigation^38^, the encoding of body posture^39^, global motion analysis^40,41^, and representations of spatial information^42^. Intriguingly, more than 50% of neurons in the RL area are multi-sensory in mice; integrating both tactile and visual sensory inputs^43^. To advance our understanding of the behavioral function of area RL/A, it will thus be important to determine whether translation- and rotation-selective neurons display multi-sensory representations of self-motion (for example, whether they encode the direction of whisker deflections). Moreover, identifying the specific projection targets of these neurons might provide insight into how sensory self-motion information feeds into, for example, neuronal circuits for movement control. We speculate that area RL/A, as defined in our experiments, may be the functional correlate of the ventral intraparietal area of the PPC in monkeys, where multi-sensory representation of self-motion is utilized for goal-directed movements^44^. Thus, an intriguing question that emerges from our results is whether responses to binocular optic flow in the PPC of monkeys rely on retinal direction selectivity, as they do in the RL/A area in mice. A first step towards addressing this would be to determine whether retinal DS cells exist in non-human primates; making it possible to define common principles of visual motion processing as well as the modifications that have occurred throughout the course of evolution.

## Acknowledgements

We thank Zoltan Raics for developing our visual stimulation system, and Bjarke Thomsen and Misugi Yonehara for technical assistance. We also thank Eric Nicholas, Ubadah Sabbagh, Thomas Wheatcroft and Lesley Anson for commenting on the manuscript. We acknowledge the following grants for financial support: Lundbeck Foundation PhD Scholarship (R230-2016-2326) to R.N.R., Velux Foundation Postdoctoral Ophthalmology Research Fellowship (27786) to A.M., Lundbeck Foundation (DANDRITE-R248-2016-2518; R252-2017-1060), Novo Nordisk Foundation (NNF15OC0017252), Carlsberg Foundation (CF17-0085), and European Research Council Starting (638730) grants to K.Y.

## Author contributions

R.N.R. and K.Y. conceived the project and designed all experiments. R.N.R. performed all viral injections and surgeries. R.N.R. performed all intrinsic signal optical imaging and two-photon calcium imaging experiments. R.N.R. and S.A. performed eye movement recording experiments. R.N.R., S.A. and A.M. analyzed the data. K.Y. provided input on all aspects of the project. R.N.R., A.M. and K.Y. wrote the manuscript.

## Methods

### Mice

All experimental procedures were approved by the Danish National Animal Experiment Committee (2020-15-0201-00452) and were performed in compliance with the Guide for the Care and Use of Laboratory Animals. Wild-type mice (C57BL/6J) were obtained from Janvier Labs. *Frmd7^tm^* mice were homozygous female or hemizygous male *Frmd7^tm1b(KOMP)Wtsi^* mice, obtained as *Frmd7^tm1a(KOMP)Wtsi^* from the Knockout Mouse Project (KOMP) Repository^24,27^: Exon 4 and the neo cassette flanked by loxP sequences were removed by crossing with female Cre-deleter *Edil3^Tg(Sox2-cre)1Amc/J^* mice (The Jackson Laboratory: stock 4783), as confirmed by PCR of genome DNA, and maintained in a C57BL/6J background. Experiments were performed on 9 male and female wild-type mice, and 8 female and male *Frmd7^tm^* mice. All mice were 12–18 weeks old during imaging experiments. Mice were kept on a reversed 12 h dark/light cycle and housed in groups of up to four littermates per cage.

### Chronic cranial windows

Mice were anaesthetized with an intraperitoneal injection of a Fentanyl (0.05 mg/kg body weight; Hameln), Midazolam (5.0 mg/kg body weight; Hameln), and Medetomidine (0.5 mg/kg body weight; Domitor, Orion) mixture. To prevent neural edema during or after surgery, dexamethasone (0.2 mg/kg body weight; Dexium, Bimeda) was injected subcutaneously. Body temperature was maintained using a feedback-controlled heating pad (ATC2000, World Precision Instruments) and eyes were protected from dehydration with eye ointment (Viscotears, Novartis). The scalp overlying the skull was removed, and a custom head-fixing imaging head-plate, with a circular 8 mm diameter opening, was mounted using a mixture of cyanoacrylate-based glue (Super Glue Precision, Loctite) and dental cement (Jet Denture Repair Powder). The center of the head-plate was positioned above V1 (stereotaxic coordinates: 2.5 mm lateral, 1 mm anterior of lambda). A 5 mm craniotomy was made in the center of the head-plate. After removing the skull flap, the cortical surface was kept moist with Ringer’s solution (in mM): 110 NaCl, 2.5 KCl, 1 CaCl_2_, 1.6 MgCl_2_, 10 glucose, and 22 NaHCO_3_. A 5 mm glass coverslip (0.15 mm thickness, Warner Instruments) was placed onto the brain to shield and gently compress the underlying cortex. The cranial window was sealed using a cyanoacrylate-based glue (Super Glue Precision, Loctite) mixed with black dental cement (Jet Denture Repair Powder mixed with iron oxide powdered pigment), to prevent light contamination from the visual display. In addition, a black Oring was mounted on top of the head-plate to further prevent any light contamination during imaging. Mice were administered subcutaneous analgesia (0.1 mg/kg body weight; Temgesic, Indivior) and returned to their home cage after anesthesia was reversed with an intraperitoneal injection of a Flumazenil (0.5 mg/kg body weight; Hameln) and Atipamezole (2.5 mg/kg body weight; Antisedan, Orion Pharma) mixture.

### Virus injections

Mice were anesthetized with an intraperitoneal injection of a Fentanyl (0.05 mg/kg body weight; Hameln), Midazolam (5.0 mg/kg body weight; Hameln), and Medetomidine (0.5 mg/kg body weight; Domitor, Orion) mixture. To prevent neural edema during or after the surgery, dexamethasone (0.2 mg/kg body weight; Dexium, Bimeda) was injected subcutaneously. Three small 0.4 mm diameter craniotomies were made and ~100–150 nL AAV2/1-Syn-GCaMP6f-WPRE (2.13 × 10^13^ vg/ml, Penn Vector Core #AV-1-PV2822) slowly injected (5 min/injection) at a depth of ~275 μm below the dura. Injections were made using a borosilicate glass micropipette (30 μm tip diameter) and a pressure injection system (Picospritzer III, Parker). The micropipette was advanced at a 20° angle relative to vertical to minimize compression of the brain. To prevent backflow during withdrawal, the micropipette was kept at the injection site for 10 min before it was slowly retracted. The skin was sutured shut and postoperative analgesia was administered subcutaneously (0.1 mg/kg body weight; Temgesic, Indivior). Mice were returned to their home cage after anesthesia was reversed with an intraperitoneal injection of a Flumazenil (0.5 mg/kg body weight; Hameln) and Atipamezole (2.5 mg/kg body weight; Antisedan, Orion Pharma) mixture.

### Intrinsic signal retinotopic mapping

Before two-photon calcium imaging, cortical visual areas of each mouse were identified by intrinsic signal optical imaging as previously described^13^. Mice were anesthetized with isoflurane (2–3% induction) and head-fixed in a custom holder. Chlorprothexine was administered intraperitoneally (2.5 mg/kg body weight; Sigma) as a sedative^33^, and isoflurane reduced to 0.5–1% during visual stimulation. Core body temperature was maintained at 37–38 °C using a feedback-controlled heating pad (ATC2000, World Precision Instruments). The stimulated contralateral eye was kept lubricated by a thin layer of silicone oil. A 2× air-objective (Olympus, 0.08 NA) was mounted on our Scientifica VivoScope, equipped with a CMOS camera (HD1-D-D1312-160-CL-12, PhotonFocus). The camera was connected to a Matrox Solios (eCL/XCL-B) frame-grabber via Camera Link. The microscope was defocused 400–600 μm down from the pial surface, where intrinsic signals were excited using a red LED (KL1600, Schott) delivered through a 610 nm long-pass filter (Chroma). Reflected light was captured through a 700 ± 50 nm band-pass filter (Chroma) positioned in front of the camera, and images were collected at 6 frames per second. The 47.65 × 26.87 cm (width × height) display was angled 30° from the midline of the mouse and the perpendicular bisector was 10 cm from the bottom of the display, centered on the display left to right, and 10 cm from the eye^13,28^. This resulted in a visual field coverage from −41.98° to 60.77° (total 102.75°) in elevation, and from −67.23° to 67.23° (total 134.46°) in azimuth. Retinotopic maps were generated by sweeping a spherically corrected (https://labrig-ger.com/blog/2012/03/06/mouse-visual-stim/) full-field bar across the display. The bar contained a flickering black-and-white checkerboard pattern on a black background. The width of the bar was 12.5° and the checkerboard square size was 25°. Each square alternated between black and white at 4 Hz. In each trial, the bar was drifted ten times in each of the four cardinal directions, moving at 8–9 °/s. Usually, two to four trials resulted in well-defined retinotopic maps. From the raw image data, we used the response time course for each pixel and computed the phase and magnitude of the Fourier transform at the visual stimulus frequency^45^. The phase maps were then converted into retinotopic coordinates from the geometry of our setup. From this, we identified visual area borders based on the visual field sign maps and superimposed those borders with the anatomical blood-vessel images to accurately localize visual cortical areas.

### Two-photon calcium imaging

Imaging was initiated two weeks after virus injections. Mice were awake during all imaging sessions as previously described^13^. To habituate mice to handling and the experimental conditions, one week after cranial window implantation, each mouse was head-fixed onto the imaging stage with its body restrained in a cylindrical cover, reducing struggling and overt body movements^13^. The habituation procedure was repeated for at least three days for each mouse at durations of 15, 30, and 60 min on days one, two, and three, respectively. At the end of each session, mice were rewarded with chocolate paste. Imaging session lasted 1–2 hours. The area targeted for two-photon imaging was localized by previous intrinsic signal optical imaging. Imaging was performed from layer 2/3, 120–275 μm below the dura, using a Scientifica VivoScope with a 7.9 kHz resonant scanner running SciScan, and with dispersion-compensated 940 nm excitation provided by a mode-locked Ti:Sapphire laser (MaiTai DeepSee, Spectra-Physics) through an Olympus 25× (1.05 NA) objective. The emitted fluorescence photons were reflected off a dichroic mirror (525/50 nm) and collected using a GaAsP photomultiplier tube (Scientifica). Clear ultrasound gel (NeurGel, Spes Medica) was used as immersion medium. To prevent light leakage from the visual stimulation, the objective was shielded with black tape, in addition to the O-ring mounted on top of the head-plate, and black cloth covered the microscope. Average excitation laser power varied from 40 to 65 mW. Images had 512 × 512 pixels, at 0.2 μm per pixel, and were acquired at 30.9 Hz using bidirectional scanning. We observed no sign of GCaMP6f bleaching during experiments. Each mouse was imaged repeatedly over the course of 2–3 weeks.

### Visual stimulus for two-photon calcium imaging

For visual stimulation during two-photon calcium imaging experiments, two 47.65 × 26.87 cm (width × height) displays were angled 30° from the midline of the mouse on the left and right side; each display subtending 115.61° in azimuth and 80.95° in elevation (Fig. 1b). The visual stimulus protocol employed was adapted from a previous study^3^. Full-field vertical sinusoidal gratings (100% contrast; spatial frequency of 0.03 cycles/°) with a spherical correction to simulate projection onto a virtual sphere moved horizontally at speeds of 10 or 40 °/s. The horizontal transition consisted of eight separate conditions (6 s each, interspersed with 4 s of gray screen between conditions): 1) left nasal, 2) left temporal, 3) right nasal, 4) right temporal, 5) contraversive, 6) ipsiversive, 7) forward, 8) backward (Supplementary Movie 1). Conditions 1–4 and were thus monocular, and conditions 5–8 binocular, simulating the rotational and translational optic flow experienced during turning and straight movements, respectively. The sequence of eight conditions was repeated in six trials. The mouse’s binocular visual field (central 40°) did not contain the visual stimulus, to ensure only stimulation of the monocular visual field^46^.

### Eye movement tracking

In a subset of experiments, we tracked eye movements in awake mice during presentation of our visual stimulus protocol (Extended Data Fig. 2). We employed an eye-tracking system developed in our laboratory and recently described in detail^47^. Briefly, a small 45° hot mirror was aligned above a CCD camera (Guppy Pro F-031, AlliedVision) lateral to the position of the mouse. The camera was positioned below the visual field. Behind the visual stimulus display, a near-infrared light source (SLS-02082-B, Mightex Systems) was angled at 45° to illuminate the recorded eye. The camera was connected to a PC via a dedicated frame grabber (FIW62, ADLINK) and images were collected at ~65 frames per second. Using the eye-tracking software, EyeLoop, images were processed, and pupil and corneal reflection coordinates were computed^47^. From these, the angular eye coordinates (x and y) were calculated^47^. Horizontal eye speed was obtained by taking the first derivative of the horizontal eye coordinates (Vx and Vy), and low pass filtering Vx and Vy with a 1 s moving average filter^25^. Saccades were identified as events with a speed > 20 °/s. Stimulus-triggered horizontal eye speed and saccade rate traces were obtained by averaging over all trials.

### Preprocessing of two-photon calcium imaging data

Imaging data were excluded from analysis if motion along the z-axis was detected. Raw two-photon imaging movies were corrected for in-plane motion using a piecewise non-rigid motion correction algorithm implemented in MATLAB (Mathworks)^48^. To detect regions of interest (ROIs) we used the MATLAB implementation of Suite2p^49^. ROIs were automatically detected using the motion-corrected frames and afterwards manually curated using the Suite2p graphical user interface. From the motion-corrected movies and detected ROIs, we extracted the fluorescence time courses within each ROI. To correct the calcium traces for contamination from the surrounding neuropil, we also extracted the fluorescence of the surrounding neuropil for each ROI. The time series of the neuropil decontaminated calcium trace, F_d_(t), was described by:

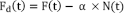

where F(t) is the somata calcium trace, N(t) is the neuropil trace, and a is the contamination factor. The contamination factor was determined for each ROI as previously^49^. Briefly, F and N traces were first low pass filtered using the 8^th^ percentile in a 180 s moving window, yielding F_s_ and N_s_, respectively. These were then used to establish F_f_(t) = F(t) — F_s_(t) and N_f_(t) = N(t) — N_s_(t). F_f_ and N_f_ were then used to determine a as previously described^49,50^. Using the neuropil decontaminated calcium trace, baseline calcium fluorescence, was computed for each stimulus condition as the mean during the pre-stimulus period ^10^. Fluorescence values were then converted to relative change compared to baseline according to: ΔF/F = (F_d_ — F)/F, where F_d_ is the instantaneous neuropil decontaminated calcium trace and F is the baseline calcium fluorescence. The mean neuronal responses were computed as the average response during the visual stimulus, and the mean and standard deviation across trials for each stimulus condition was computed for each neuron. To identify neurons for further in-depth analysis we used three inclusion criteria: 1) Neurons were defined as visually responsive if their mean AF/F to the preferred stimulus condition exceeded 10%; 2) A response reliability index, δ, was computed for each neuron as:

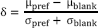

where μ_max_ and σ_max_ are the mean and standard deviations of the response to the preferred stimulus condition respectively, and μ_blank_ and σ_blank_ are the mean and standard deviations of the response to a blank stimulus respectively^10^. Neurons with δ exceeding 0.6 were defined as reliable; and 3) A signal-to-noise ratio (SNR) was computed for each neuron as:

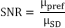

where μ_pre_ is the mean of the response to the preferred stimulus condition and μ_SD_ is the mean of the standard deviation of the fluorescence trace during the baseline period (0.5 s before stimulus onset) for each trial^51^. Neurons with SNR exceeding 0.5 were defined as robustly responding. Only neurons that fulfilled all inclusion criteria at both stimulus speeds were included for further analysis procedures.

### Response profile classification

In order to classify the response of individual neurons into separate functional groups, representing distinct response profiles, we employed regression analysis similar to previously described^3^. First, we summarized the response of each neuron by a tuning curve, including the mean ΔF/F for each of the eight stimulus conditions. We compiled this tuning curve for both stimulus speeds, and we determined the speed in which the highest mean AF/F was evoked; noted as the preferred speed of the neuron. By considering the response selectivity of a neuron to the eight stimulus conditions, we assumed that the response profile regressors could be described by an indicator function, *R*, as follows:

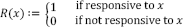

where *x* is the stimulus condition, and 2^8^ (i.e. 256) possible regressors exist for *R* (Fig. 1g). These 256 regressors correspond to the possible response combinations from the monocular and binocular stimulations in the nasal and temporal directions. For each neuron we then computed the linear Pearson’s correlation for its tuning curve at the preferred speed against each of the 256 regressors and determined the regressor with the highest correlation. All neuronal tuning curves had high correlation with its assigned response regressor (mean correlation coefficient, 0.91 ± 0.05, *n* = 26712 neurons from 17 mice). The response regressors were functionally described using a MATLAB implementation of the Quine and McCluskey algorithm (https://www.mathworks.com/matlabcentral/fileexchange/37118-mintruthtable-tt-flags), in which the Boolean functions were minimized to find the logical function for each response profile that use only a small number of logical operations^3^. Here, we focused on the simple (MoDS: regressors IDs, 75, 43, 61, 56; BiDS: 194, 206, 193, 211; and non-DS: 234, 247, 256), binocular-suppressed (DS: 6, 7, 8, 9; and non-DS: 24, 37), and translation-selective or rotation-selective (FT: 32, 17, 80, 3; BT: 25, 20, 68, 2; CR: 33, 22, 85, 4; and IR: 28, 19, 67, 5) response classes (Fig. 1h and Extended Data Fig. 4). The simple and translation- or rotation-selective response classes are responsive to both monocular and binocular motion stimulation, and these were identified and described in detail previously^3^. In this work, we identified the binocular-suppressed response class, characterized by only responding to monocular motion stimulation, in a DS or non-DS manner. The binocular-suppressed functional groups (regressor IDs) were described by the following Boolean logical operations:

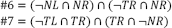

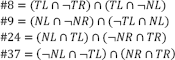

where # is the identity of the regressors, and *N* and *T* are nasal and temporal motion directions, respectively, and *L* and *R* are stimulation of the left and right eyes, respectively, and ¬ is a logical “NOT” gate operator.

### Comparison of response classes and functional groups among visual cortical areas

To examine similarities and disparities in response class distributions across visual areas in wild-type and *Frmd7^tm^* mice, we performed a hierarchical clustering analysis. For this, we used the mean proportion of the nine functional groups (simple: MoDS, BiDS, and non-DS; binocular-suppressed: DS and non-DS; and translation- or rotation-selective: FT, BT, IR, and CR) to create a monocular and binocular motion flow “fingerprint” for each visual area in wild-type and *Frmd7^tm^* mice. To create a hierarchical cluster tree, we used the *linkage* function in MATLAB, and visualized the result in a dendrogram (Fig. 4a). For quantifying similarities and disparities across visual areas within wild-type and *Frmd7^tm^* mice (Fig. 4b), we computed Pearson’s correlation coefficients using the motion flow fingerprint of each visual area. To quantify the proportions of monocular versus binocular functional groups within each visual cortical area, we computed a binocular optic flow index (BOFI). For this, we determined the proportion of monocular driven (simple MoDS and non-DS, and binocular-suppressed DS and non-DS) and binocular driven (simple BiDS, and FT, BT, CR, and IR) functional groups, and computed the BOFI as:

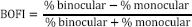

with a BOFI of 1 indicating that only binocular driven functional groups are represented, while a BOFI of −1 indicates only monocular driven groups are represented.

### Quantification and statistical analysis

To statistically evaluate populational differences in functional groups between wild-type and *Frmd7^tm^* mice, we performed a two-way ANOVA test followed by post hoc comparisons using the two-sided Mann-Whitney *U* test. To compare Pearson’s correlation coefficients obtained from two independent samples, i.e. wild-type and *Frmd7^tm^* mice, we used the Fischer’s r-to-z transformation and obtained the corresponding two-sided *P* value. Center and spread values are reported as mean ± s.e.m. We used no statistical methods to plan sample sizes but used sample sizes similar to those frequently used in the field^10,13,17^. Exact *n* (i.e. number of animals and neurons) is included in the Result section and Supplementary Table 1. *P* < 0.05 was considered statistically significant, where *P < 0.05 and * *P < 0.01. Statistical analyses were carried out in MATLAB.

---

**Extended Data Fig. 1.**
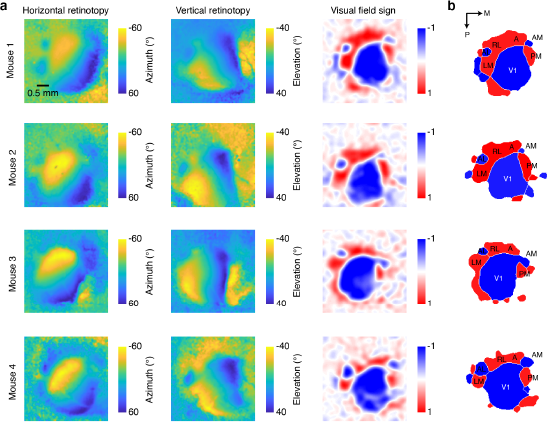
Identification of visual cortical areas using intrinsic signal optical imaging. **a**, Maps of horizontal (left) and vertical retinotopy (middle), and the corresponding visual field sign map (right) from four example mice. **b**, Thresholded visual field sign patches showing the location of primary visual cortex (V1), and the higher visual areas: lateromedial (LM), anterolateral (AL), rostrolateral (RL), anterior (A), anteromedial (AM), and posteromedial (PM). Coordinates indicate posterior (P) and medial (M) directions.

**Extended Data Fig. 2.**
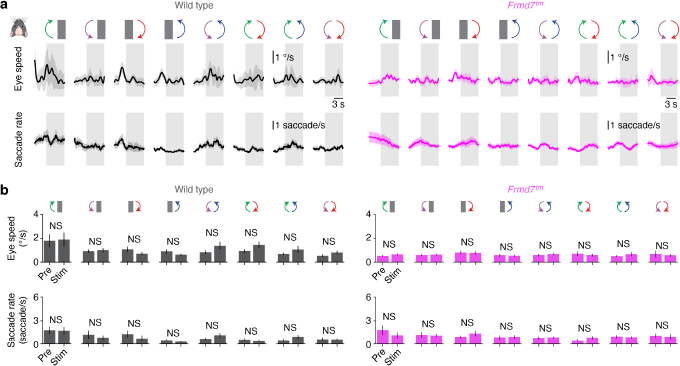
Eye movements in awake wild-type and *Frmd7^tm^* mice during visual stimulus protocol. **a**, Trial-averaged horizontal eye speed and saccade rate time courses recorded in wild-type mice (left; *n* = 9 recordings) and *Frmd7^tm^* mice (right; *n* = 9 recordings) in response to the monocular and binocular horizontal motion conditions presented at 10 °/s. Error bars are mean ± s.e.m. **b**, Quantification of mean horizontal eye speed (upper) and mean saccade rate (lower) before and during visual stimulation in wild-type (left) and *Frmd7^tm^* mice (right) (NS, not significant, *P* ≥ 0.05; Wilcoxon signed-rank test; *n* = 9 recordings from 3 wild type mice and 9 recordings from 3 *Frmd7^tm^* mice). Error bars are mean ± s.e.m.

**Extended Data Fig. 3.**
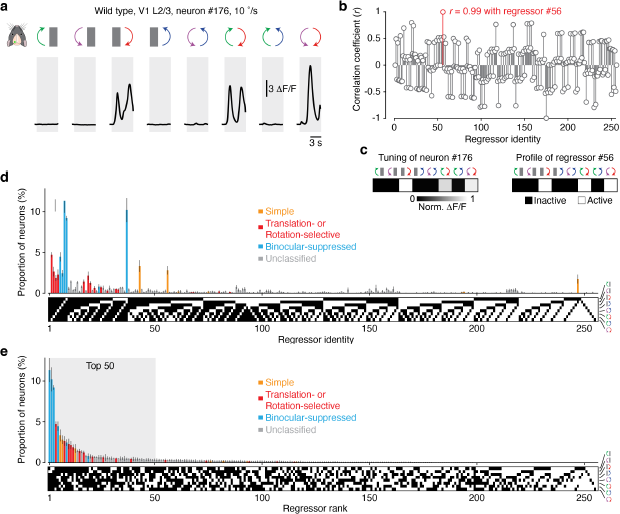
Regression analysis for classifying individual neurons to discrete response types. **a**, Trial-averaged fluorescence intensity (ΔF/F) time course for example layer 2/3 V1 neuron (#176) from a wild-type mouse in response to the monocular and binocular motion conditions at 10 °/s. **b**, The tuning curve at the preferred speed was correlated to each of the 256 regressors, yielding a correlation profile. Correlation coefficients were calculated as Pearson’s *r*. Neuron #176 showed the highest correlation with regressor #56 (*r* = 0.99) and was thus assigned to this response type. **c**, Tuning profile of neuron #176 and response profile of regressor #56. **d**, Distribution of all reliably responsive V1 neurons from wild-type mice *(n* = 3010 neurons from 4 mice) grouped according to the 256 regressors and response class (simple, translation- or rotation-selective, binocular-suppressed, and unclassified). Error bars are mean ± s.e.m. **e**, Distribution from (**d**) ranked according to regressor frequency. The shaded region depicts the 50 most abundant regressors (as shown in Fig. 2).

**Extended Data Fig. 4.**
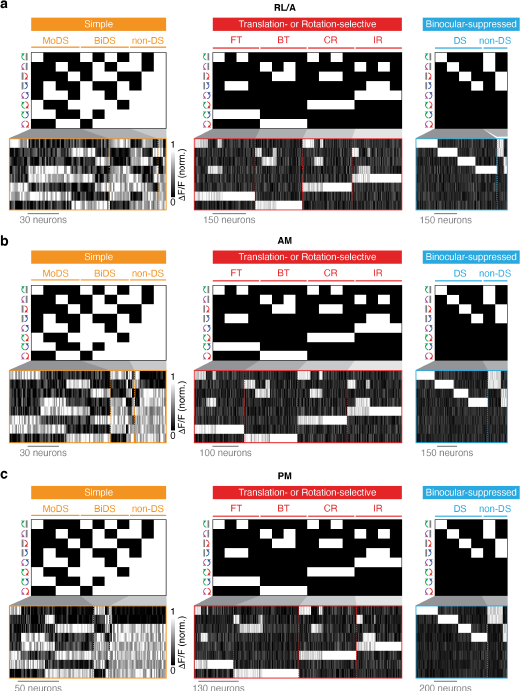
Tuning of higher visual area neurons assigned to functional groups. **a,b,c** Regressor profiles and tuning of RL/A (**a**), AM (**b**) and PM (**c**) neurons assigned to functional groups within simple, translation- or rotation-selective, and binocular-suppressed response classes. MoDS: monocular DS; BiDS: binocular DS; FT: forward translational; BT: backward translational; CR: contraversive rotational; IR: ipsiversive rotational.

**Supplementary Table 1.** Numbers of neurons sampled by visual cortical area and genetics. Total neurons: total number (*n*) of neurons recorded in wild-type (WT) and *Frmd7^tm^* mice experiments for each visual cortical area. Consistently responsive: number (*n*) and percent of total of neurons that met the inclusion criteria for responsiveness (ΔF/F > 10%), reliability (δ > 0.5), and signal-to-noise (SNR > 0.5) and were included for regressor correlation analysis. Animals: number (*n*) of WT and *Frmd7^tm^* mice that data were collected from for each area.

| Area | Total neurons |  | Consistently responsive |  | Animals |  |
| --- | --- | --- | --- | --- | --- | --- |
|  | WT: *n* | *Frmd7<sup>tm</sup>*: *n* | WT: *n* (%) | *Frmd7<sup>tm</sup>*: *n* (%) | WT: *n* | *Frmd7<sup>tm</sup>*: *n* |
| V1 | 5748 | 5534 | 3010 (52%) | 2925 (53%) | 4 | 4 |
| RL/A | 6563 | 6746 | 4165 (63%) | 3125 (46%) | 5 | 5 |
| AM | 6664 | 6919 | 4006 (60%) | 3375 (49%) | 5 | 5 |
| PM | 5419 | 5868 | 3059 (56%) | 3047 (52%) | 4 | 4 |

## Reference

1. Krapp, H. G., Hengstenberg, R. & Egelhaaf, M. Binocular Contributions to Optic Flow Processing in the Fly Visual System. J. Neurophysiol. 85, 724–734 (2001).

2. Farrow, K., Haag, J. & Borst, A. Nonlinear, binocular interactions underlying flow field selectivity of a motion-sensitive neuron. Nat. Neurosci. 9, 1312–1320 (2006).

3. Kubo, F. et al. Functional Architecture of an Optic Flow-Responsive Area that Drives Horizontal Eye Movements in Zebrafish. Neuron 81, 1344–1359 (2014).

4. Naumann, E. A. et al. From Whole-Brain Data to Functional Circuit Models: The Zebrafish Optomotor Response. Cell 167, 947-960.e20 (2016).

5. Wylie, D. R. W., Bischof, W. F. & Frost, B. J. Common reference frame for neural coding of translational and rotational optic flow. Nature 392, 278–282 (1998).

6. Simpson, J. I., Leonard, C. S. & Soodak, R. E. The accessory optic system of rabbit. II. Spatial organization of direction selectivity. J. Neurophysiol. 60, 2055–2072 (1988).

7. Sunkara, A., DeAngelis, G. C. & Angelaki, D. E. Joint representation of translational and rotational components of optic flow in parietal cortex. Proc. Natl. Acad. Sci. U. S. A. 113, 5077–82 (2016).

8. Duffy, C. J. & Wurtz, R. H. Sensitivity of MST neurons to optic flow stimuli. I. A continuum of response selectivity to large-field stimuli. J. Neurophysiol. 65, 1329–1345 (1991).

9. Tanaka, K. & Saito, H. A. Analysis of motion of the visual field by direction, expansion/contraction, and rotation cells clustered in the dorsal part of the medial superior temporal area of the macaque monkey. J. Neurophysiol. 62, 626–641 (1989).

10. Marshel, J. H., Garrett, M. E., Nauhaus, I. & Callaway, E. M. Functional specialization of seven mouse visual cortical areas. Neuron 72, 1040–1054 (2011).

11. Zhuang, J. et al. An extended retinotopic map of mouse cortex. Elife 6, (2017).

12. Cruz-Martín, A. et al. A dedicated circuit links direction-selective retinal ganglion cells to the primary visual cortex. Nature 507, 358–61 (2014).

13. Rasmussen, R., Matsumoto, A., Dahlstrup Sietam, M. & Yonehara, K. A segregated cortical stream for retinal direction selectivity. Nat. Commun. 11, 831 (2020).

14. Glickfeld, L. L., Andermann, M. L., Bonin, V. & Reid, R. C. Cortico-cortical projections in mouse visual cortex are functionally target specific. Nat. Neurosci. 16, 219–26 (2013).

15. Lyamzin, D. & Benucci, A. The mouse posterior parietal cortex: Anatomy and functions. Neurosci. Res. 140, 14–22 (2019).

16. Hovde, K., Gianatti, M., Witter, M. P. & Whitlock, J. R. Architecture and organization of mouse posterior parietal cortex relative to extrastriate areas. Eur. J. Neurosci. 49, 1313–1329 (2019).

17. Minderer, M., Brown, K. D. & Harvey, C. D. The Spatial Structure of Neural Encoding in Mouse Posterior Cortex during Navigation. Neuron 102, 232-248.e11 (2019).

18. Gilissen, S. R. J., Farrow, K., Bonin, V. & Arckens, L. Reconsidering the border between the visual and posterior parietal cortex of mice. bioRxiv 2020.03.24.005462 (2020) doi:10.1101/2020.03.24.005462.

19. Dhande, O. S. & Huberman, A. D. Retinal ganglion cell maps in the brain: implications for visual processing. Curr. Opin. Neurobiol. 24, 133–142 (2014).

20. Borst, A. & Euler, T. Seeing Things in Motion: Models, Circuits, and Mechanisms. Neuron 71, 974–994 (2011).

21. Rasmussen, R. & Yonehara, K. Contributions of Retinal Direction Selectivity to Central Visual Processing. Curr. Biol. 30, R897–R903 (2020).

22. Wei, W. & Feller, M. B. Organization and development of direction-selective circuits in the retina. Trends in Neurosciences vol. 34 638–645 (2011).

23. Yonehara, K. et al. Identification of Retinal Ganglion Cells and Their Projections Involved in Central Transmission of Information about Upward and Downward Image Motion. PLoS One 4, e4320 (2009).

24. Hillier, D. et al. Causal evidence for retina-dependent and -independent visual motion computations in mouse cortex. Nat. Neurosci. 20, 960–968 (2017).

25. Macé, É. et al. Whole-Brain Functional Ultrasound Imaging Reveals Brain Modules for Visuomotor Integration. Neuron 100, 1241-1251.e7 (2018).

26. Sabbah, S. et al. A retinal code for motion along the gravitational and body axes. Nature 546, 492–497 (2017).

27. Yonehara, K. et al. Congenital Nystagmus Gene FRMD7 Is Necessary for Establishing a Neuronal Circuit Asymmetry for Direction Selectivity. Neuron 89, 177–193 (2016).

28. Juavinett, A. L., Nauhaus, I., Garrett, M. E., Zhuang, J. & Callaway, E. M. Automated identification of mouse visual areas with intrinsic signal imaging. Nat. Protoc. 12, 32–43 (2016).

29. Andermann, M. L., Kerlin, A. M., Roumis, D. K., Glickfeld, L. L. & Reid, R. C. Functional Specialization of Mouse Higher Visual Cortical Areas. Neuron 72, 1025–1039 (2011).

30. Kretschmer, F., Tariq, M., Chatila, W., Wu, B. & Badea, T. C. Comparison of optomotor and optokinetic reflexes in mice. J. Neurophysiol. 118, 300–316 (2017).

31. Dougherty, K., Cox, M. A., Westerberg, J. A. & Maier, A. Binocular Modulation of Monocular V1 Neurons. Curr. Biol. 29, 381–391 (2019).

32. Wang, Q., Sporns, O. & Burkhalter, A. Network analysis of corticocortical connections reveals ventral and dorsal processing streams in mouse visual cortex. J. Neurosci. 32, 4386–99 (2012).

33. Smith, I. T., Townsend, L. B., Huh, R., Zhu, H. & Smith, S. L. Stream-dependent development of higher visual cortical areas. Nat. Neurosci. 20, 200–208 (2017).

34. Ramachandra, V., Pawlak, V., Wallace, D. J. & Kerr, J. N. D. Impact of visual callosal pathway is dependent upon ipsilateral thalamus. Nat. Commun. 11, (2020).

35. Wertz, A. et al. Single-cell-initiated monosynapic tracing reveals layer-specific cortical network modules. Science (80-.). 349, 70–74 (2015).

36. Yonehara, K. et al. The First Stage of Cardinal Direction Selectivity Is Localized to the Dendrites of Retinal Ganglion Cells. Neuron 79, 1078–1085 (2013).

37. Lien, A. D. & Scanziani, M. Cortical direction selectivity emerges at convergence of thalamic synapses. Nature 558, 80–86 (2018).

38. Gold, J. I. & Shadlen, M. N. The neural basis of decision making. Annual Review of Neuroscience vol. 30 535–574 (2007).

39. Mimica, B., Dunn, B. A., Tombaz, T., Srikanth Bojja, V. P. T. N. & Whitlock, J. R. Efficient cortical coding of 3D posture in freely behaving rats. Science (80-.). 362, 584–589 (2018).

40. Juavinett, A. L. & Callaway, E. M. Pattern and Component Motion Responses in Mouse Visual Cortical Areas. Curr. Biol. 25, 1–6 (2015).

41. Rasmussen, R. & Yonehara, K. Circuit Mechanisms Governing Local vs. Global Motion Processing in Mouse Visual Cortex. Front. Neural Circuits 11, (2017).

42. Save, E. & Poucet, B. Role of the parietal cortex in long-term representation of spatial information in the rat. Neurobiol. Learn. Mem. 91, 172–178 (2009).

43. Olcese, U., Iurilli, G. & Medini, P. Cellular and synaptic architecture of multisensory integration in the mouse neocortex. Neuron 79, 579–593 (2013).

44. Bremmer, F. Navigation in space - The role of the macaque ventral intraparietal area. J. Physiol. 566, 29–35 (2005).

